# Microglia and meningeal macrophages depletion delay the onset of experimental autoimmune encephalomyelitis

**DOI:** 10.1101/2022.06.10.495612

**Authors:** Alejandro Montilla, Alazne Zabala, Marco Er-Lukowiak, Björn Rissiek, Tim Magnus, Noelia Rodriguez-Iglesias, Amanda Sierra, Carlos Matute, María Domercq

**Author notes:** Correspondence should be addressed to: María Domercq, Dpt. Neuroscience, Universidad del País Vasco, E-48940 Leioa, Spain.

## Abstract

In multiple sclerosis and the experimental autoimmune encephalomyelitis (EAE) model, both resident microglia and infiltrating macrophages contribute to demyelination as well as spontaneous remyelination. Nevertheless, the specific roles of microglia versus macrophages are unknown. We investigated the influence of microglia in EAE using the colony stimulating factor 1 receptor (CSF-1R) inhibitor, PLX5622, to deplete microglial population and *Ccr2*^RFP/+^*fms*^EGFP/+^ mice, to distinguish peripheral macrophages and microglia. PLX5622 treatment depleted microglia and meningeal macrophages, and provoked a massive infiltration of CCR2^+^ macrophages into demyelinating lesions and spinal cord parenchyma. PLX5622 treatment did not alter EAE chronic phase. In contrast, microglia and meningeal macrophages depletion reduced the expression of CD80 co-stimulatory molecule in dendritic and myeloid cells and reduced T cell reactivation and proliferation in the spinal cord parenchyma, inducing a significant delay in EAE onset. Altogether, these data points to a specific role of CNS microglia meningeal macrophages in antigen presentation and T cell reactivation at initial stages of the EAE model.

## INTRODUCTION

Multiple sclerosis (MS) is a chronic inflammatory disease of the brain and spinal cord leading to demyelination and neurodegeneration. These outcomes are caused by immune cell infiltration across the blood–brain barrier, promoting detrimental effects such as inflammation, gliosis, and neuroaxonal degeneration ^1^. In experimental autoimmune encephalomyelitis (EAE), an MS mouse model, myelin-specific T cells are peripherally activated in secondary organs, such as lymph nodes, and migrate towards the CNS interfaces to get reactivated by antigen presenting cells (APCs) and clonally expand to effectively carry out their actions ^2, 3^. Beside T cells, myeloid cells such as resident microglia, CNS-associated macrophages (CAMs, found in interfaces such as the perivascular space or the meninges) or infiltrating monocytes are associated to the development of the MS/EAE pathology. All these myeloid populations are known to participate in both beneficial and detrimental processes regarding the development of the pathology. Reactive microglia and monocyte-derived macrophages are thought to contribute to neurodegeneration as their number correlates with the extent of axonal damage in MS lesions ^4–7^. The response of microglia/macrophages may represent one of the initial steps in EAE pathogenesis, preceding and possibly triggering T cell development and infiltration of blood-derived cells ^8–12^. However, other studies indicate that microglia/macrophages activation counteracts pathological processes by providing neurotrophic and immunosuppressive factors and by promoting recovery ^13–15^. These dichotomic behaviour could be explained on the basis of different activation states ^16, 17^, as modulation of microglia/macrophage activation determine EAE outcome and potentiate recovery ^14, 18^. On the other hand, it has been previously hypothesized that macrophages and microglia could play different roles along the pathology. Monocyte-derived macrophages associate with nodes of Ranvier and contributes to demyelination, whereas microglia appear to clear myelin debris ^11^. Accordingly, microglia during EAE display a weakly immune-activated phenotype whereas infiltrated macrophages are highly immune reactive ^19^. Nevertheless, the specific differences in the roles of resident microglia and infiltrating monocytes is still unknown, due to the difficulty in distinguishing these two populations ^20^.

Microglia and CAMs, but not peripheral monocytes, arise from early progenitors in the embryonic yolk sac, that migrate and colonize the CNS, in a process controlled by the colony stimulating factor-1 receptor (CSF-1R) and its ligands ^21, 22^. CSF-1R inhibition has been proposed as specific for central nervous system (CNS) microglia depletion without significant effect on peripheral immune cells (1–4) and as a strategy to unveil the role of these cells. Treatment with PLX5622, a selective inhibitor of CSF-1R, results in microglial population depletion as it has been defined as an experimental strategy to characterize the role of microglia in physiology or pathology. In this study, we caused microglial depletion using PLX5622, administered two weeks prior to the EAE induction and along the course of the EAE, to specifically study microglial implication at all disease stages.

## MATERIALS AND METHODS

### Animals

All experiments were performed according to the procedures approved by the Ethics Committee of the University of the Basque Country (UPV/EHU). Animals were handled in accordance with the European Communities Council Directive. All possible efforts were made to minimize animal suffering and the number of animals used. To generate *Ccr2*^RFP/+^*fms*^EGFP/+^ double transgenic mice, *Ccr2*^RFP/RFP^ and *fms*^EGFP/EGFP^ mice were crossed and first-generation littermates were used. Both transgenic lines are on a C57BL/6 genetic background.

### Microglial depletion

To deplete microglia *in vivo*, C57BL/6 and *Ccr2*^RFP/+^*fms*^EGFP/+^ mice were fed with 1200 ppm PLX5622 (Plexxikon Inc.) *ad libitum*. Respective control animals received standard chow instead. Mice were fed for at least 21 days prior to further experimental procedures to ensure maximal microglial depletion.

### EAE induction

EAE was induced in 8- to 10-week-old female C57BL/6 and *Ccr2*^RFP/+^*fms*^EGFP/+^ mice. Mice were immunized with 200 µg of myelin oligodendrocyte glycoprotein 35-55 (MOG_35–55_; MEVGWYRPFSRVVHLYRNGK) in incomplete Freund’s adjuvant (IFA; Sigma) supplemented with 8 mg/ml *Mycobacterium tuberculosis* H37Ra (Fisher). Pertussis toxin (500 ng; Sigma) was injected intraperitoneally on the day of immunization and 2 days later. Neurological score was assessed daily and ranged from 0 to 8 as previously described ^23^. Depending on the experiment, mice were euthanized at different phases of the model, which are defined as pre-onset (prior to the appearance of the first symptoms; day post-immunization (dpi) 8-9), onset (mice show first symptomatology; dpi 10-14) and chronic phase (mice have stabilized neurological score, dpi 30-35).

After EAE, mice are euthanized and spinal cords were removed. The different regions were differentially processed; cervical and thoracic regions were flash frozen for subsequent biochemical analysis, while lumbar region was drop-fixed in 4% paraformaldehyde (PFA). Peripheral immune-related organs such as spleen or lymph nodes were also frozen for qPCR analysis. For flow cytometry analysis, peripheral blood was also extracted right after the euthanasia and collected in lithium-heparin tubes to avoid coagulation.

### Immunohistochemistry

In order to perform histological analysis on the spinal cords after EAE, tissues were fixed in 4% PFA for 3-4 hours, and subsequently transferred to 15% sucrose for at least 2 days. Then, lumbar regions were frozen in 15% sucrose - 7% gelatine solution in PBS, and 12-mm coronal sections were obtained. Primary antibodies used for immunohistochemistry were: mouse anti-myelin basic protein (MBP) (1:1000; Covance), rabbit anti-MBP (1:200; Millipore), rabbit anti-Iba1 (1:500; Wako Chemicals), mouse anti-GFAP (1:40; Millipore), rabbit anti-mannose receptor (1:200; Abcam), rat anti-CD3 (1:50; Bio-Rad), rat anti-CD45R (1:200; BD Bioscience), rat anti-Ly6G (1:100; BioLegend), mouse anti-SMI32 (1:1000; Covance), rabbit anti-Ki67 (1:500, Vector Laboratories), mouse anti-CD31 (1:100; Santa Cruz), rat anti-GFP (1:200;) and rabbit anti-dsRed (1:600; Clontech). The anti-GFP and anti-dsRed antibodies were used to amplify the intrinsic fluorescent signals in the *Ccr2*^RFP/+^*fms*^EGFP/+^ mice. These primary antibodies were subsequently detected by incubation with appropriate Alexa Fluor 488 or 594 conjugated goat antibodies (1:250; Invitrogen). Images were acquired using a Leica TCS STED SP8 confocal microscope or a Zeiss LSM800 confocal microscope with the same settings for all samples within one experimental group. All the image analysis was performed with the ImageJ software (National Institutes of Health).

### Evaluation of blood-brain barrier disruption

To analyze whether the blood-brain barrier permeabilization was different in the experimental groups, animals were injected intraperitoneally with 2% Evans Blue (EB) in saline (200 µL), and let it spread throughout the body for 1 hour. Spinal cord tissue was fixed and processed as described above, and EB staining in blood vessels was evaluated by immunohistochemistry.

### Flow cytometry

For the fluorescence-activated single cell (FACS) analysis, immune cells were isolated from spleen, spinal cord and peripheral blood. Specifically, spleens were mashed through 70 µm cell strainers using a syringe piston. Alternatively, spinals cords were processed by both enzymatical and mechanical digestion. During the protocol, erythrocytes were lysed using an ACK lysis buffer (155 mM NH_4_Cl, 10 mM KHCO_3_, 0.1 mM EDTA, pH 7.2). Single cell suspensions were stained with fluorochrome-conjugated monoclonal antibodies in buffer containing 1 mM EDTA (Sigma) and 0.1 % bovine serum albumin (Sigma). Regarding the immune cell profiling, the antibodies used were the following: CD3-PE/Cy7, CD11b-Bv510, CD8-Bv650, CD4-Bv785, CD80-PE, CD11c-Bv605, CD45-APC/Cy7, Ly6G-AF700, TCRgd-perCP/Cy5.5, CD86-Bv421, P2Y12-APC and MHCII-FITC. All these antibodies were used in a 1:100 concentration, and were acquired from BioLegend. Cells were analyzed using a BD FACSCelesta, and all the data was analyzed with FlowJo software (BD Bioscience).

Regarding the measurement of resident microglia and invading macrophages populations in spinal cord during EAE chronic phase, we used CD11b-FITC (1:200; BioLegend) and CD45-PE (1:100; BioLegend) as antibodies, identifying microglia as the CD11b^+^/CD45^low^ population, and infiltrating monocytes as the CD11b^+^/CD45^hi^ population. These analyses were performed using a BD FACSJazz cell sorter and analyser.

### Serum cytokines quantification

In parallel to the FACS analysis, part of the serum from both control and PLX5622-treated animals’ blood was separated in order to measure different pro- and anti-inflammatory cytokines. Specifically, the levels of IFN-γ, TNF-α, IL-2, IL-6, IL-17A and IL-22 were measured using a LEGENDplex™ Mouse Th Cytokine Panel (BioLegend), according to the manufacturer’s instructions.

### Quantitative RT-PCR

Total RNA from EAE lumbar spinal cords, spleens and lymph nodes was isolated using TRIzol (Invitrogen) according to the manufacturer’s instructions. Afterwards, 2 µg of this RNA was used to perform a retrotranscription protocol, using SuperScript III Reverse Transcriptase (200 U/μL; Invitrogen) and random hexamers as primers (Promega).

Real-time quantitative PCRs (qPCRs) were conducted in a Bio-Rad Laboratories CFX96 real-time PCR detection system, as previously described ^24^. The reactions were performed using SYBR Green as the DNA-binding dye and specific primers for different T cell subtypes (Table I). The primers were designed using Primer Express Software (Applied Biosystems) at exon junctions to avoid genomic DNA amplification. The cycling conditions comprised 3 min of polymerase activation at 95°C and 40 cycles consisting of 10 s at 95°C and 30 s at 60°C. The amount of cDNA was quantified using a standard curve from a pool of cDNA obtained from the different conditions of the experiment. Finally, the results were normalized using a normalization factor based on the geometric mean of housekeeping genes (Table I) obtained for each condition using the geNorm v3.5 software ^25^.

**Table I.**
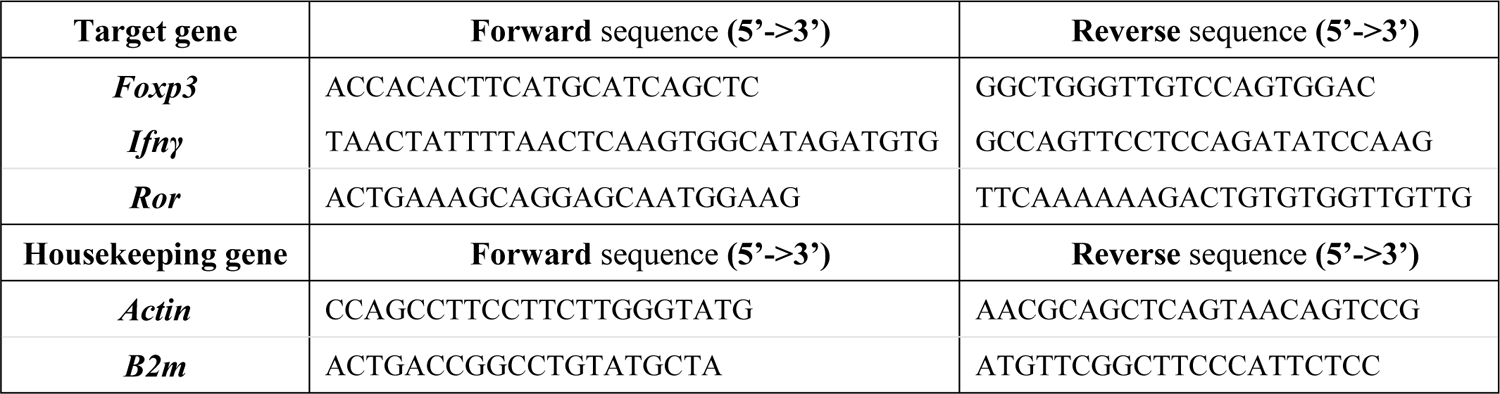
Sequences of primers used for qPCRs

### Statistical analysis

Data are presented as mean ± SEM with sample size and number of repeats indicated in the figure legends. All the statistical analyses were performed with GraphPad Prism 8.0 (GraphPad Software). Specifically, every comparison between two groups was analyzed using unpaired Student’s two-tailed t test. Comparisons among multiple groups were analyzed by one-way analysis of variance (ANOVA) followed by Bonferroni’s multiple comparison tests. Statistical significance was considered when p < 0,05.

## RESULTS

### PLX5622 treatment causes a delay in the EAE onset and a massive infiltration of macrophages

Following EAE induction, CNS-resident microglia and invading macrophages contribute to disease pathogenesis, influencing both progression and recovery of the disease ^26^. To address the role of microglial cells at all EAE stages, we used the CSF-1R specific inhibitor PLX5622 (administered at 1200 ppm in chow) to delete microglia. We eliminated 90% of microglia throughout the lumbar spinal cord after 3 weeks of treatment, in a similar proportion of that achieved in other studies ^27, 28^ (Fig. 1A). We further characterized whether PLX5622 treatment affected CNS-associated macrophages, which could be distinguished from microglia by the expression of mannose receptor C-type1 (MRC1, also known as CD206) ^29^. In control mice, PLX5622 treatment also reduced the number of Iba1^+^ CD206^+^ cells at the meninges (Fig. 1B), demonstrating that PLX5622 also depleted meningeal macrophages.

**Figure 1.**
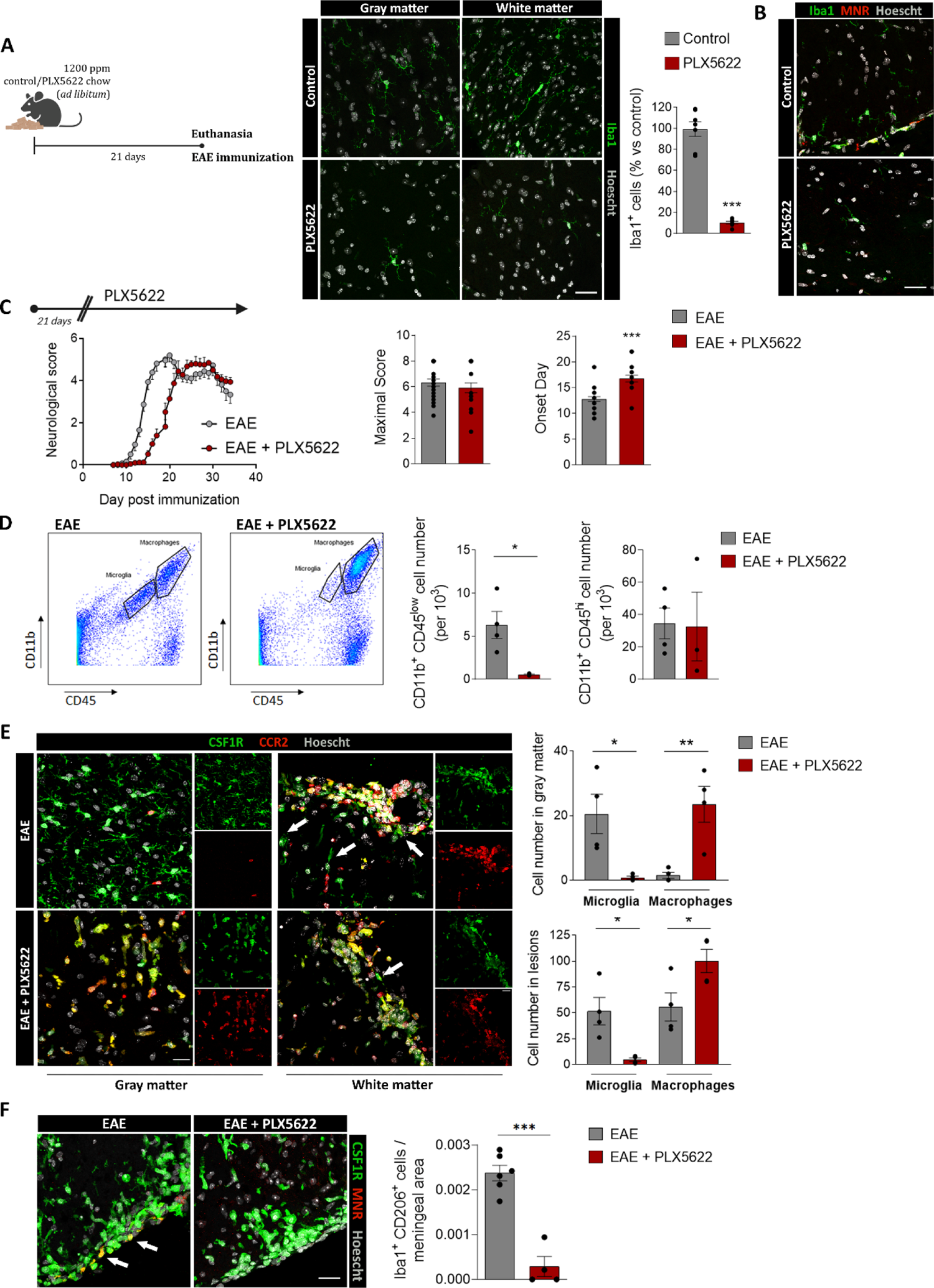
PLX5622 microglial depletion provokes a delay in the onset of EAE and massive infiltration of macrophages. ***(A)*** (Left) Scheme showing the paradigm of microglial depletion with PLX5622. (Right) Representative images showing microglial (Iba1^+^ cells) depletion in both white and gray matter of healthy spinal cord. Histogram shows the loss of microglia after PLX5622 treatment, in comparison to control mice. Scale bar = 25 µm. ***(B)*** Representative images of Iba1^+^ MNR^+^ meningeal macrophages in the spinal cord of control mice and mice after complete PLX5622 treatment. Scale bar = 25 µm***. (C)*** (Left) Neurological score of control and PLX5622-treated mice, (n = 20-25 mice from 3 independent EAE experiments). (Right) Histograms showing the maximal score reached by every animal, as well as the onset day of clinical signs. ***(D)*** Plots depicting the strategy to distinguish resident microglia (CD11b^+^/CD45^low^) from invading macrophages (CD11b^+^/CD45^hi^) in EAE spinal cords from control and PLX5622-treated mice. Histograms show the quantification of both populations, in relation to the total number of analyzed cells in the samples (n = 4 mice per group). ***(E)*** Representative images of *fms*^+^ microglia (green, arrows) and *Ccr2*^+^ macrophages (red) in the gray and white matter of control and PLX5622-treated Ccr2^RFP/+^fms^EGFP/+^ mice at EAE chronic phase. Histograms at the right show the quantification of microglia (fms^+^ CCR2^-^ cells) and macrophages (CCR2^+^ cells) in each region (n = 3). Scale bar = 20 µm. ***(F)*** Representative images of *fms^+^* MNR^+^ meningeal macrophages (arrows) in the spinal cord of control and PLX5622-treated EAE mice. Scale bar = 25 µm. Histogram shows the number of meningeal macrophages in relation to the area of the meninges analyzed. Data are presented as means ± SEM. *p < 0.05, ***p < 0.001

Induction of EAE after treatment with PLX5622 for 3 weeks delayed the onset of symptoms but not the subsequent disease course (Fig. 1C). We did not observe differences in the maximal neurological score reached at the peak of the disease, nor in the recovery capacity during the chronic phase of EAE (Fig. 1C). Then, we assessed the populations of microglia (CD11b^+^ CD45^low^) and invading macrophages (CD11b^+^ CD45^hi^) at the end of the disease by FACS, and we observed that the ablation of microglia was maintained throughout the EAE (Fig. 1D).

To understand the dynamics of CNS microglia and macrophages, as well as invading monocytes, we induced EAE in control and PLX5622-treated *Ccr2*^RFP/+^*fms*^GFP/+^. These mice allowed us to distinguish resident microglia and CAMs (*Ccr*2^-^*fms*^+^) vs infiltrating macrophages (*Ccr*2^+^ *fms*^-^) on spinal cord sections. Of note, infiltrated macrophages (*Ccr*2^+^) started to express *fms-EGFP*, so we also considered *Ccr*2^+^*fms*^+^ cells as infiltrating macrophages. Indeed, the CNS environment induces a microglial-like phenotype in myeloid populations ^30^. PLX5622 chronic treatment in these mice (as described before) induced a massive reduction of *Ccr*2^-^ *fms*^+^ microglia cells as well as *Ccr*2^-^ *fms*^+^ CD206^+^ meningeal macrophages at EAE chronic phase (Fig. 1E, F). In contrast, we observed a of peripheral macrophages into demyelinated lesions in white matter in PLX5622-treated mice (Fig. 1E). In addition, macrophages penetrated into the non-damaged white matter and even into gray matter in PLX5622-treated mice (Fig. 1E). These results demonstrate that resident microglia and menigeal macrophages limit the massive entry of macrophages and their dispersion through the CNS parenchyma in response to pathological conditions. Interestingly, despite the massive infiltration of macrophages, microglia and menigeal macrophages depletion did not trigger an exacerbated progression of EAE, nor did it cause a failure in the recovery phase, as described earlier ^9^. Together, these results indicate that microglial and menigeal macrophages ablation provokes a compensatory mechanism based on a robust infiltration of peripheral macrophages along with a delayed disease onset.

### Microglia and menigeal macrophages depletion does not alter EAE chronic pathophysiology

Microglial function during EAE is commonly associated with its capacity to phagocytose myelin debris, promoting recovery and regeneration in the chronic phase of the disease. We further assessed whether depletion of microglia affects neuroinflammation and demyelinated lesions at this stage using immunohistochemistry. The extent of the lesioned area was determined by the accumulation of infiltrating cells, and by the loss or damage of myelin (characterized by high MBP immunoreactivity).

We observed no differences in the white matter area affected by EAE between control and PLX5622-treated mice (Fig. 2A). Demyelination in EAE is mediated by an immune response, mainly based on T cell activity but also on B cells infiltration ^31^. Because of that, we analyzed the presence of both populations in the lesions and found no differences in meningeal and infiltrating accumulation of both cell types (Fig. 2B). Lastly, we observed no alteration in astrogliosis neither in the lesions nor in the surrounding parenchyma (Fig. 2B).

**Figure 2.**
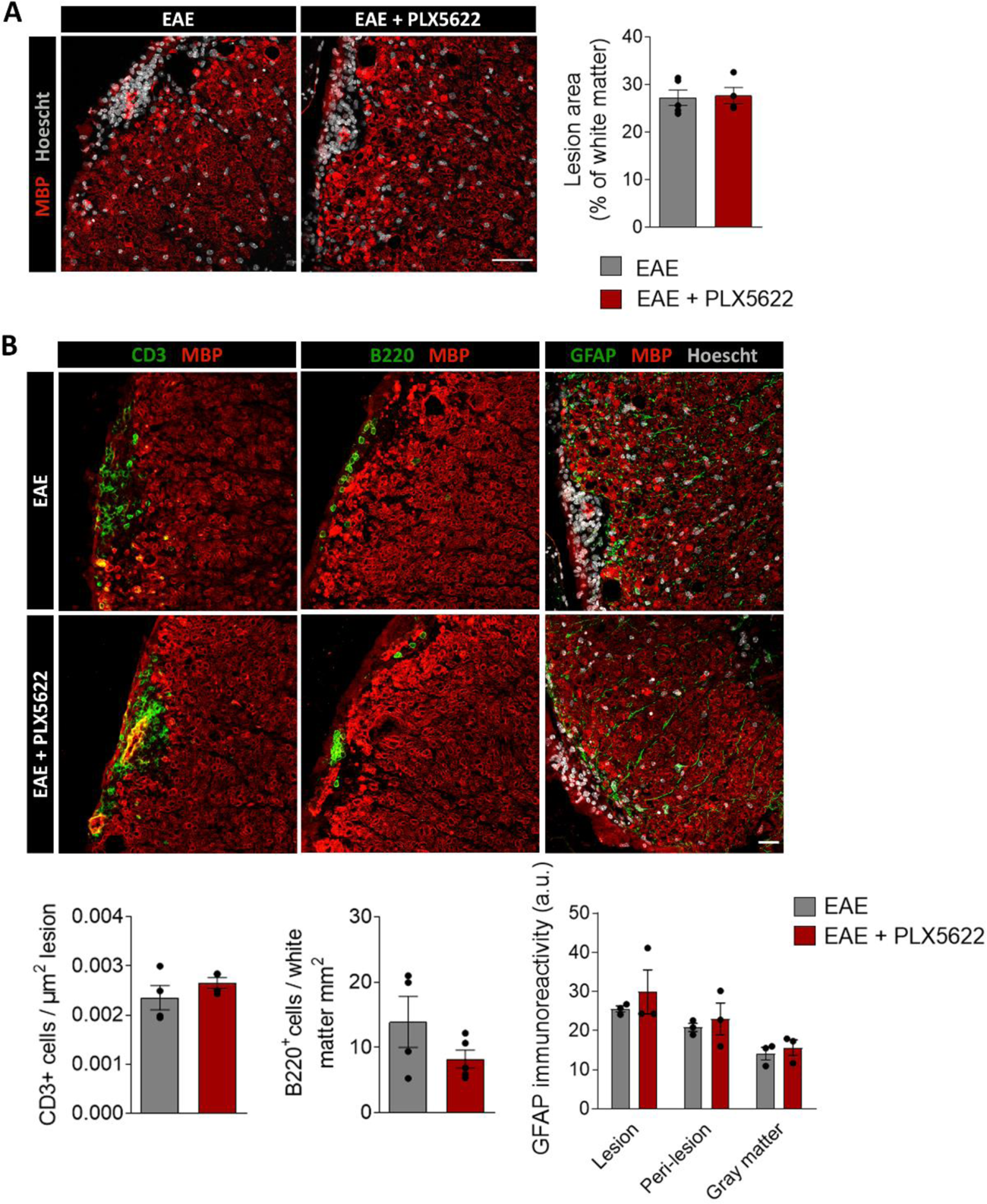
PLX5622 microglial depletion does not alter the EAE chronic phase. *(**A)*** Representative images of EAE lesions in the lumbar spinal cord of control and PLX5622-treated mice, at 35 days post-immunization. Histogram shows the percentage of lesioned white matter versus total white matter (n = 5). Scale bar = 50 µm. ***(B)*** Representative images showing the accumulation of CD3^+^ T cells and B220^+^ B cells in EAE lesions as well as the astrogliosis in control and PLX5622-treated at EAE chronic phase (35 days after post-immunization). Scale bar = 25 µm. Histograms show the number of cells normalized to lesion area or total white matter for T and B cells, respectively (n = 5), and astrocyte immunoreactivity, indicative of astrogliosis (n = 3). Data are presented as means ± SEM.

In sum, these findings show that the course of the chronic phase of EAE was not altered in mice with depleted microglia. As we detected similar partial remission of the symptoms in both experimental groups, this suggests that microglia are not necessary for an EAE better outcome or a more efficient remyelinating process during the disease.

### Peripheral immune priming is not altered in EAE after microglial and menigeal depletion with PLX5622

As previously described, EAE initial stages primarily imply T cell activation in peripheral lymphoid organs, such as the spleen or lymph nodes, against myelin-specific peptides ^2^. Since microglial ablation delayed EAE onset, we hypothesized that microglia could modulate immune priming or immune infiltration. In order to corroborate these hypotheses, we first assessed whether microglia could influence peripheral early responses right after immunization, and prior to the appearance of first motor deficits.

We immunized mice and euthanized them at the pre-onset phase of EAE (dpi 8-9; no clinical signs). At this timepoint, we performed flow cytometry analysis of the immune cell populations in spleen and peripheral blood (FACS gating specified in Fig. 3A). Microglial depletion by inhibition of CSF-1R did not alter the proportion of CD4^+^ T cells, CD8+ T cells, γδ T cells, neutrophils and macrophages neither in the spleen nor in the blood (Fig. 3B, C).

**Figure 3.**
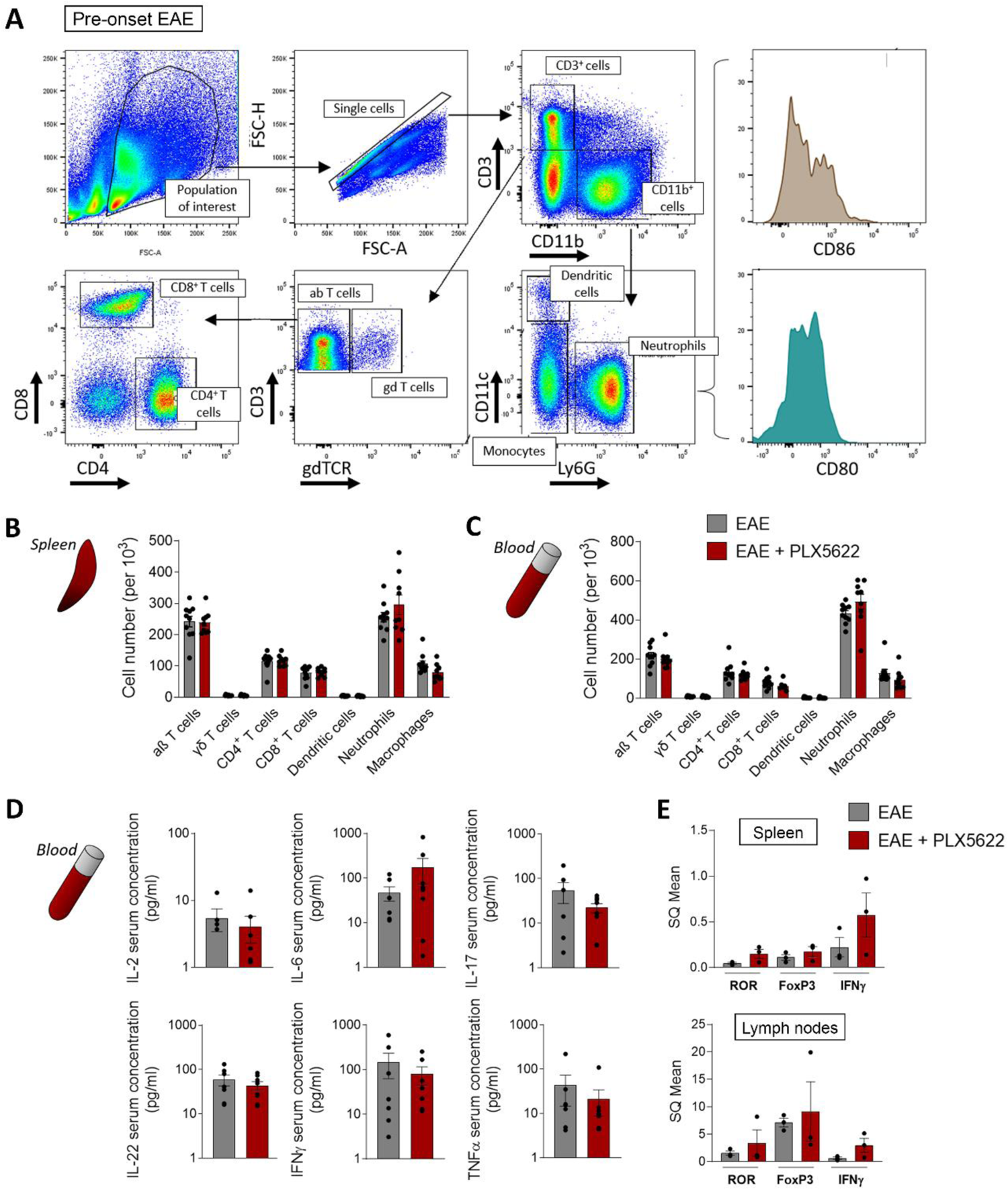
PLX5622 did not alter peripheral immune priming after immunization. ***(A)*** Flow cytometry gating strategy for analysis of immune populations in the spleen and peripheral blood of mice at EAE pre-onset (dpi 8-9). ***(B, C)*** Quantification of αβ T cells (CD3^+^ γδTCR^-^), γδ T cells (CD3^+^ γδTCR^+^), CD4 T cells (CD3^+^ CD4^+^) CD8 T cells (CD3^+^ CD8^+^), dendritic cells (CD11b^+^ CD11c^+^), neutrophils (CD11b^+^ Ly6G+), and macrophages (CD11b^+^ CD11c^-^ Ly6G^-^) populations in spleen (B) and blood (C), in relation to the total CD45^+^ single cells analyzed at this stage (n = 10). ***(D)*** Concentration of cytokines in blood serum from control and PLX5622-treated mice at EAE pre-onset stage (n = 7). ***(E)*** Relative mRNA expression of *Ror*, *Foxp3* and *Ifnγ* in spleen and lymph nodes of both mice groups, at EAE pre-onset stage (n = 3). Data are presented as means ± SEM.

Taking advantage of the blood sampling, we analyzed the concentration of pro- and anti-inflammatory cytokines in the serum of both animal groups, using a bead-based immunoassay. Specifically, we measured the levels of innate and adaptive immune cytokines such as IL-2, IL-6, IL-17, IL-22, IFNγ and TNFα, all associated with EAE pathology ^32^. Microglial ablation did not lead to any difference in the cytokine profiles in the serum at EAE pre-onset (Fig. 3D). Moreover, as EAE model is based predominantly in a CD4+ T cell response, we assessed their phenotypes by qPCR analysis in spleen and lymph nodes, both tissues where T cells are early activated. We measured the levels of mRNA expression for forkhead box protein P3 (*FoxP*3), retinoic acid-related orphan receptor (*Ror*), transcription factors specifying Treg and Th17 activity respectively ^33, 34^, as well as *Ifn*γ, signature cytokine for Th1 cells ^35^. We did not find differences in the expression of any of these markers between control and PLX5622-treated mice at EAE pre-onset (Fig. 3E).

All these results suggest that microglial and CAMs ablation does not provoke significant alterations in the primary, peripheral immune priming after EAE induction. Thus, the delay in the onset of the symptoms might be associated with an effect of PLX5622 within the CNS environment.

### Spinal cord early EAE affection is not altered after microglial and meningeal depletion

As the peripheral response was not altered in mice treated with the CSF-1R inhibitor, we next analyzed the spinal cord of both control and PLX5622-treated mice at the pre-onset stage of EAE, in order to look for early signs of alteration that would lead to symptoms delay.

We assessed immune cell infiltration into the spinal cord by flow cytometry profiling at EAE pre-onset (gating strategy specified in Fig. 4A). Aside from the expected significant reduction in the number of microglial cells in the PLX5622-treated animals, we did not identify any other differences in immune populations (Fig. 4B). Moreover, analysis of first-arriving CD4^+^ T cells’ profiles in spinal cord by qPCR did not show any differences between both experimental groups (Fig. 4C). Since alterations in BBB permeability are a key initiating factor promoting the infiltration of myeloid and lymphoid cells to the CNS parenchyma in MS and EAE ^36^, we subsequently analyzed the state of BBB integrity by histological analysis of EB in spinal cord vascularity. No differences were observed in the extravasation of EB between control and PLX5622-treated mice at EAE pre-onset, with all the staining remaining inside the CD31^+^ blood vessels in both groups (Fig. 4D).

**Figure 4.**
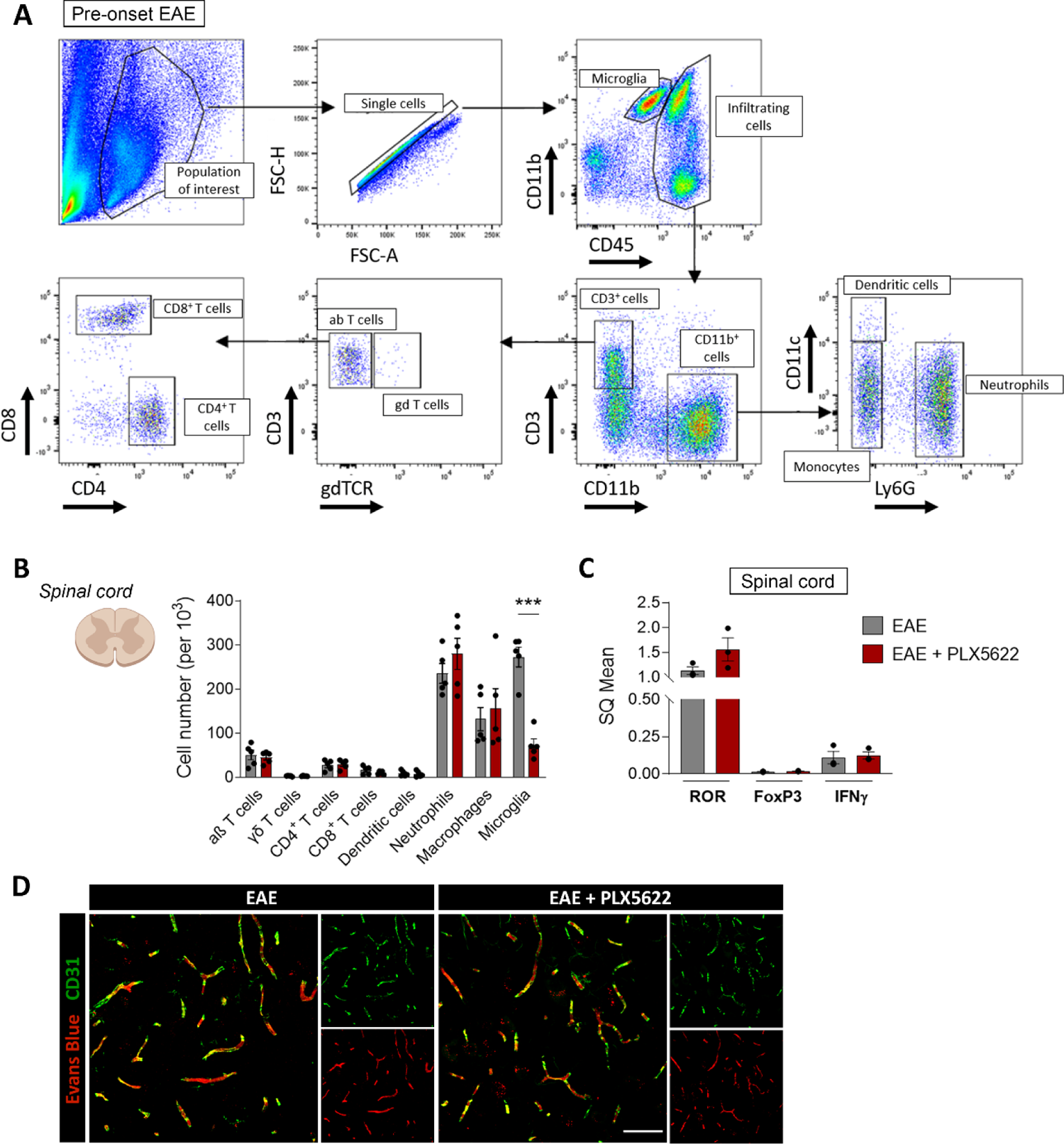
PLX5622 did not alter early, CNS-related events at the pre-onset EAE. ***(A)*** Flow cytometry gating strategy for analysis of immune populations from the spinal cord of mice at EAE pre-onset (dpi 8-9). ***(B)*** Quantification of the same populations as in the previous figure plus microglia (CD11b^+^ CD45^low^) in spinal cord, in relation to the total number of analyzed cells, at this stage (n = 5). ***(C)*** Relative mRNA expression of *Ror*, *Foxp3* and *Ifng* in the spinal cord of both mice groups, at EAE pre-onset stage (n = 3). ***(D)*** Representative images showing Evans Blue staining restricted to CD31^+^ blood vessels, showing a lack of BBB disruption in the spinal cord of both control and PLX5622-treated mice. Scale bar = 50 µm. Data are presented as means ± SEM. ***p < 0.001.

These results highlight that CSF-1R inhibition does not significantly alter immune priming on the periphery at the pre-onset stage of the disease. Thus, delayed symptomatology is probably linked to a later alteration, most likely affecting events occurring within the CNS.

### Microglia and/or meningeal macrophages are key in the immune response in the CNS parenchyma

The efficient activation of T lymphocytes after arriving to the CNS parenchyma is a critical requirement for the induction of neuroinflammation and associated EAE pathology. Because of this, and given the delay in the appearance of motor alterations in microglial-depleted mice, we next analyzed whether microglia ablation altered the immune response at EAE onset in the CNS parenchyma. We induced EAE in control and PLX5622-treated mice and euthanized them at dpi 12-14, when motor symptoms became evident.

At this timepoint, we observed a consistent difference in the degree of pathology development (Fig. 5A). Indeed, control mice were starting to show the first motor deficiencies while PLX5622-treated mice did not. We performed flow cytometry analysis of the immune cell populations in peripheral tissues (spleen and blood); the gating strategy was carried out as previously described (Fig. 3A). These flow cytometry experiments showed no alteration in the immune cell populations in spleen and blood after PLX5622 treatment (Fig. 5B). However, in the spinal cord we detected an increased proportion of neutrophils as well as a lesser percentage of macrophages and CD4^+^ T cells in PLX5622-treated animals (Fig. 5C; gating strategy described in Fig. 4A). Although the proportion of the major immune populations was not affected in microglia and menigeal macrophages depleted mice, the total number of immune cells infiltrated into the CNS parenchyma was massively reduced in PLX5622-treated mice, as revealed by immunohistochemistry (Fig. 5D). Indeed, control mice presented high levels of infiltration of T cells, B cells and neutrophils, as assessed by immunostaining of CD3, B220 and Ly6G respectively, in correlation with the neurological score (Fig. 5D).

**Figure 5.**
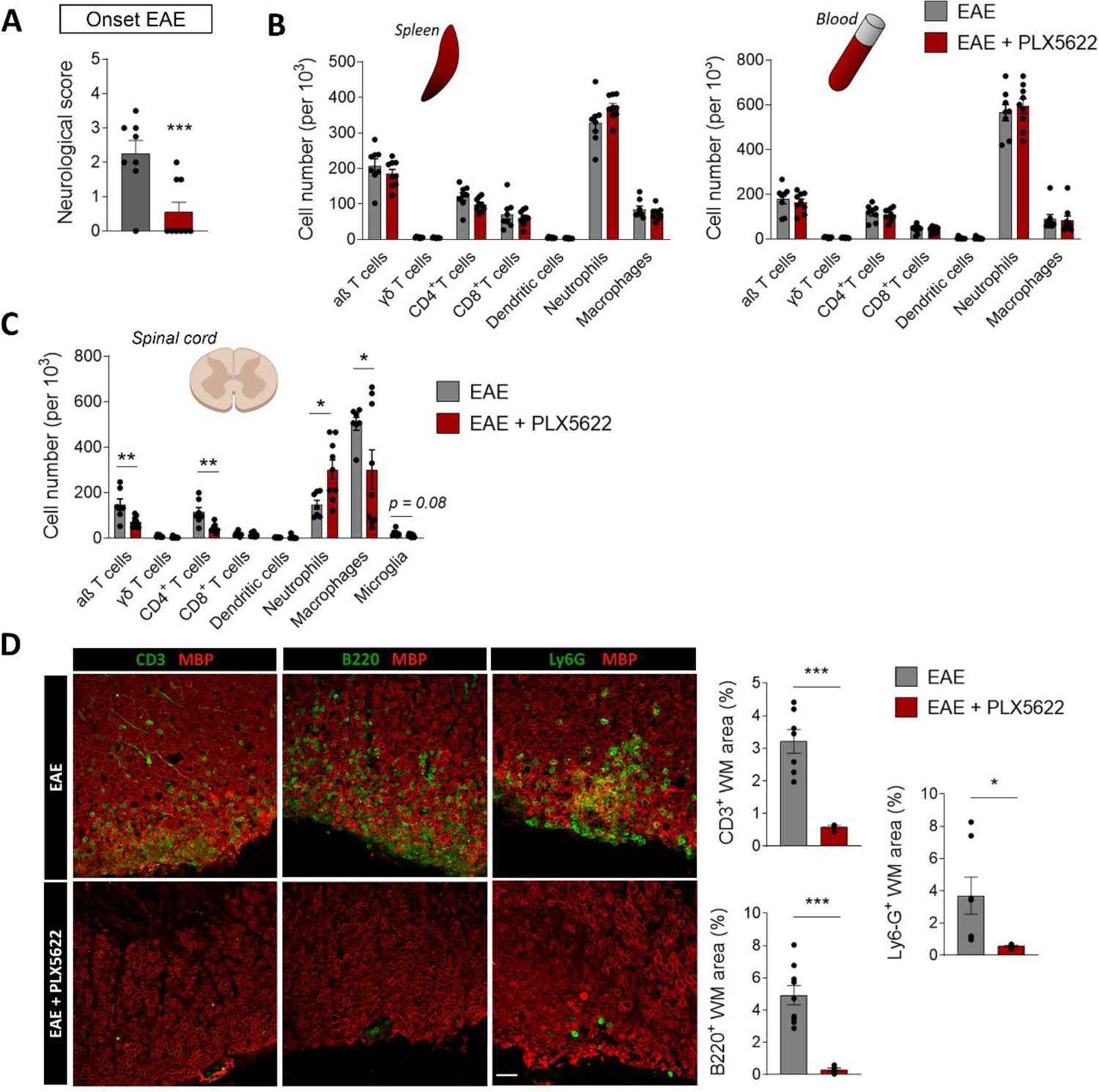
PLX5622 provoked a delay in the infiltration of immune cells towards CNS parenchyma. **(A)** Histogram showing the neurological score in control and PLX5622-treated mice at 12-14dpi. **(B)** Quantification of immune populations in spleen and peripheral blood, in relation to the total number of analyzed cells, at this timepoint (n = 10). Gating strategy is specified in Fig. 3A. **(C)** Quantification of immune populations in spinal cord, in relation to the total number of analyzed cells, at EAE onset (n = 10). Gating strategy is specified in Fig. 12A. **(D)** (Left) Representative images of CD3^+^, B220^+^ and Ly6G^+^ cells infiltration into spinal cord parenchyma. (Right) Histograms show the area occupied by these cells, in relation to the whole white matter area, in control and PLX5622-treated mice (n= ??). Scale bar = 30 µm. **(D)** Quantification of immune populations in spinal cord, in relation to the total number of analyzed cells, at EAE onset (n = 10). Gating strategy is specified in Fig. 12A. Data are presented as means ± SEM. *p < 0.05, **p<0.005, ***p < 0.001.

Alternatively, PLX5622-treated mice showed very limited infiltration of these cells, and mostly restricted to the meninges and blood vessels (Fig. 5D). These results suggest that microglia and meningeal depletion somehow provokes a delay in the accumulation of immune cells in the CNS parenchyma, and this effect is directly linked to the delay in the appearance of the motor deficits.

### Microglia and meningeal macrophages elimination reduced the expression of antigen presenting proteins and T cell reactivation in EAE onset

The accumulation of T cells in the CNS tissue during neuroinflammation is commonly preceded by a reactivation step carried out by APCs. As we observed an alteration in this accumulation, we assessed the antigen presentation process both at the pre-onset and onset stages of EAE by FACS analysis. Specifically, we measured the expression of the B7 co-stimulatory molecules (CD80 and CD86), which participate along with MHC-II in this mechanism in diverse APCs ^37^. At the pre-onset phase of EAE (dpi 8-9), we hardly found any significant differences regarding CD80 and CD86 intensity in the cell types analyzed (microglia, DCs and macrophages) in PLX5622-treated mice (Fig. 6A), except for a decrease in CD86 intensity in spleen macrophages. However, at EAE onset we detected that the expression of these co-stimulatory molecules was altered specifically in the spinal cord, but not in spleen or blood cells (dpi 12-14). The expression of CD80 was decreased in DCs and infiltrating macrophages in PLX5622-treated mice (Fig. 6B). Likewise, CD86 intensity was also reduced in macrophages of PLX5622-treated mice (Fig. 6B). These findings suggest a role of microglia and meningeal macrophages in modulating antigen presentation, not only as an intrinsic mechanism, but also affecting other APC cells and their function.

**Figure 6.**
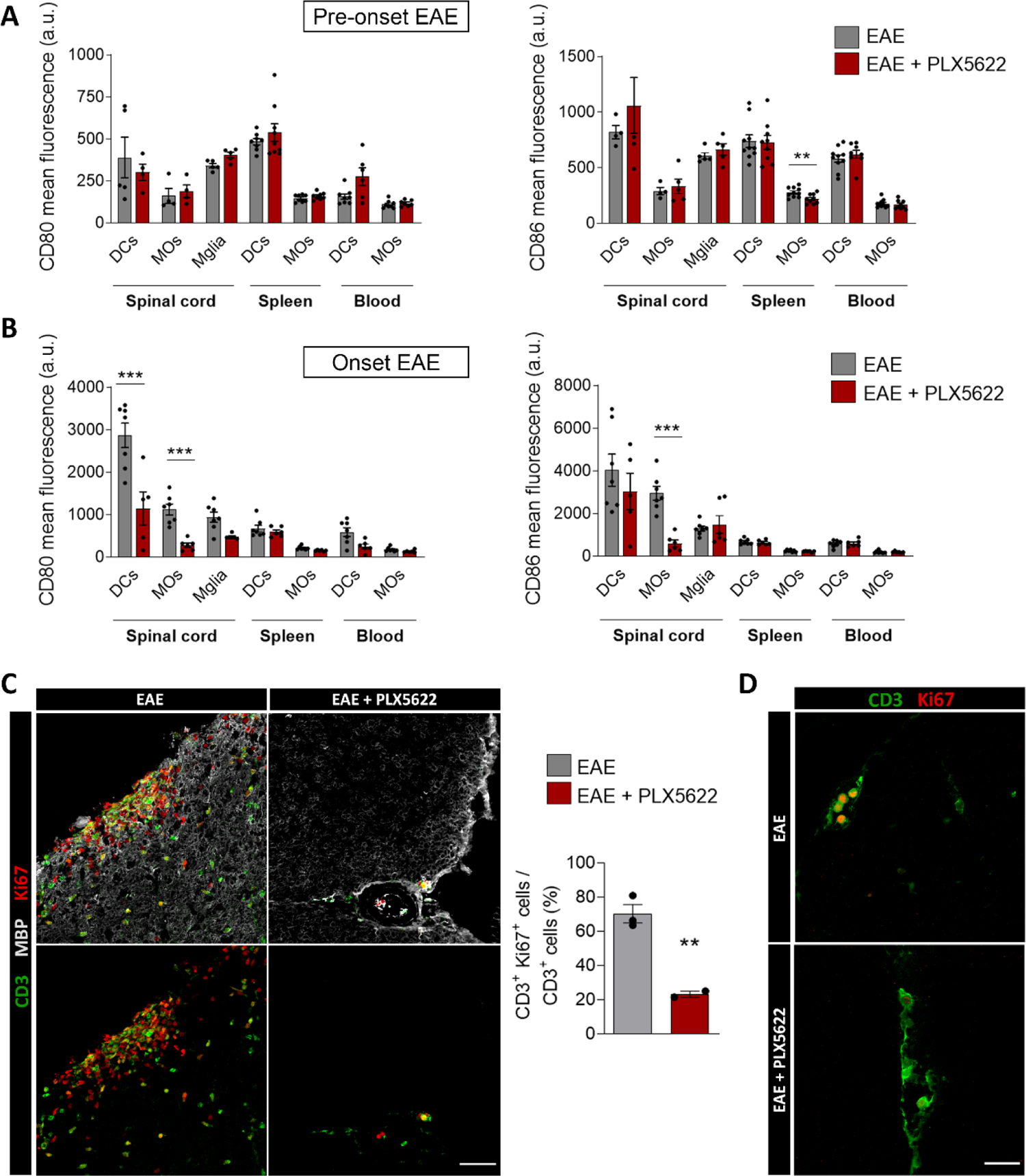
Microglial depletion with PLX5622 alters antigen presentation and T cell reactivation in the spinal cord. ***(A, B)*** Histograms showing CD80 and CD86 fluorescence intensity in different APCs (dendritic cells (DCs), macrophages (MOs) and microglia (Mglia)) and in diverse tissues, as analyzed by flow cytometry analysis, at the pre-onset stage of EAE (**A**; dpi 8-9) (n = 5 spinal cord samples, n = 10 spleen and blood samples), and at the onset stage of EAE (**B**; dpi 12-14) (n = 10). ***(C)*** Representative images showing CD3^+^ lymphocytes and Ki67 proliferative-associated expression in spinal cord, at dpi 10-12 after EAE immunization in control and PLX5622-treated mice. Histogram show the proportion of Ki67^+^ CD3^+^ T cells in relation to the total number of CD3^+^ T cells (n = 3). Scale bar = 25 µm. ***(D)*** Representative images showing the early Ki67^+^ CD3^+^ proliferating lymphocytes in the spinal cord at EAE onset (dpi 10). Scale bar = 30 µm. Data are presented as means ± SEM. **p < 0.005, ***p < 0.001.

As these observations suggest that antigen presentation was altered in spinal cord after microglial and meningeal macrophages depletion, and this process is needed for T clonal expansion in CNS parenchyma, we assessed whether PLX5622 affected T cell proliferation rate using antibodies to Ki67, a marker of proliferation. Control EAE mice showed a massive proliferation of immune cells inside the lesions, whereas proliferation was limited in PLX5622-treated mice at EAE onset (dpi 12-14; Fig. 6C). In particular, the number of Ki67^+^ CD3^+^ T cells was significantly reduced, suggesting that T cell proliferation and therefore their clonal expansion was delayed in microglia and meningeal macrophages depleted mice (Fig. 6C). Of note, this effect was also observed in the first T cells arriving to the CNS after peripheral priming (Fig. 6D).

Taken together, the results described in this work showed that microglia and meningeal macrophages limit the dispersion of CNS-infiltrated macrophages into the CNS parenchyma. In their absence, macrophages are able to colonize the CNS parenchyma and contribute to both EAE development as well as the recovery/remyelinating mechanisms occurring in the chronic phase. On the other hand, microglia and meningeal macrophages are important to EAE development as their deletion delays immune cell activation responsible for the onset of the disease, by altering the antigen presentation in the brain.

## DISCUSSION

Microglial cells participate in neurodegenerative pathologies development. Thus, the analysis of their specific role in these processes has lately emerged as an important focus of research, seeking for therapeutic approaches for diseases like MS ^38^. In this study, we used the CSF-1R antagonist, PLX5622, in order to assess the effect of microglia depletion in EAE development. However, we cannot exclude that the effect observed in EAE development after PLX5622 treatment could be partly due to the elimination of meningeal macrophages. We demonstrated that microglia and meningeal macrophages interfere with peripheral macrophage dynamics by controlling their entry and migration. The massive infiltration of macrophages in the absence of microglia and meningeal macrophages does not significantly affect neurological damage at EAE peak or recovery in EAE chronic phase. However, microglia and meningeal macrophages ablation delays EAE onset demonstrating a specific role of these cells in antigen presentation and T cell proliferation at early stages of EAE.

PLX5622 has largely been assumed to be microglia-specific but more recent studies described alterations in myeloid and lymphoid populations in peripheral tissues (Lei et al., 2020; Spiteri et al., 2022). In our study we did not find any alteration in the immune populations in spleen and blood nor in the expression of antigen presenting proteins after PLX5622 treatment, a fact suggesting that PLX5622 did not affect directly immune cells, including peripheral monocytes. While tissue macrophages and circulating monocytes express CSF-1R, their survival is not only depending on CSF1/CSF1R signaling but relies also on CCL2/CCR2 signaling ^40^ which is not present in microglia. In contrast to infiltrating and peripheral macrophages, we found that CAMs, in particular meningeal macrophages, were also depleted after PLX5622 treatment. In accordance, perivascular macrophages survival is also dependent on CSF1/CSF1R signaling, as PLX5622 treatment induced a 60% reduction in their number ^41^.

We showed that microglial and meningeal macrophages depletion provoke a massive infiltration of CCR2^+^ peripheral macrophages during EAE progression in comparison to control mice, which could constitute a compensatory mechanism. Importantly, infiltrating macrophages in PLX5622-treated mice colonize CNS parenchyma including both white and grey matter of the spinal cord, and their location is not limited to demyelinated lesions as in control mice. These data suggest that meningeal macrophages and microglia control the dispersion of macrophages throughout the CNS parenchyma. In accordance, microglia surround and confine infiltrating macrophages into the CNS parenchyma, limiting macrophage dispersion after LPC-induced demyelination ^42^. In addition, we detected a higher accumulation of macrophages in demyelinated lesions after PLX5622 treatment, suggesting that microglia and meningeal macrophages also control the infiltration, proliferation or survival of macrophages into the CNS parenchyma. In accordance, previous data using parabiosis in the EAE model demonstrated that peripheral macrophage infiltration is preceded by microglia cell death ^9^. Thus, microglia could potentially help to limit peripheral CNS inflammation and to maintain the “CNS immune-privileged” status.

PLX5622 treatment induced a consistent delay in the appearance of the first EAE symptoms. This suggests that microglia and meningeal macrophages play a role in the effector stage of the disease model. Indeed, inhibition of chemokine receptor-dependent recruitment of monocytes to the CNS blocked EAE progression, not EAE onset, suggesting that microglia is essential for EAE onset whereas macrophages contribute to EAE progression ^9^. Specifically, we have seen that microglia and meningeal macrophages ablation caused an alteration in antigen presentation to the first T cells arriving to the CNS, both in dendritic cells and infiltrated macrophages. Frequently, DCs have been identified as the main APCs during EAE ^20, 43, 44^. Interestingly, we showed that microglial and meningeal macrophages ablation reduced the expression of co-stimulatory molecules (CD80 and CD86) in all antigen presenting cells, including DC cells, in the spinal cord but not periphery, excluding the possibility of a direct effect of PLX5622 treatment. This suggests microglia and/or meningeal macrophages early activation after EAE induction potentially promote antigen presentation capacity in other cell types. This result is in accordance with those obtained in a virus model, in which PLX5622 also alters T cell local reactivation in CNS decreasing B7 co-stimulatory signals in CD11c^+^ cells ^45^. Altogether, these data highlight the relevance of microglia and meningeal macrophages in orchestrating the CNS immune response.

Numerous studies determine that microglia activation occurs during the onset and peak of EAE and its activation is necessary to EAE development. Microglia control T cell encephalogenicity through the release of IL23, specifically the P40 subunit and its deletion in microglia cells suppressed EAE by shifting T cell response towards a Th2 rather than Th1 ^46^. Another important signal for microglia activation is the TGFβ-activated kinase (TAK1). Thus, microglia selective ablation of TAK1 blocks its activation, the release of pro-inflammatory mediators such as IL1β and CCL2, and the immune cell infiltration, suppressing completely EAE ^10^. Microglial paralysis provoked by the treatment of ganciclovir (GCV) in CD11b-HSVTK transgenic mice led to a repression of clinical EAE ^8^. Overall, our data, in accordance with previous studies, indicate that microglia and its persistent and overt activation has a detrimental role in CNS autoimmunity onset, and preventing or suppressing this process may be therapeutic.

Targeting CSF-1R has also been tried in different EAE models and at different time windows leading to diverse outcomes. Inhibiting CSF-1R with PLX5622 after EAE onset attenuated EAE pathology and promoted recovery ^47^. However, blocking CSF-1R with PLX3397 at later stages in EAE exacerbated neuroinflammation and neurological damage in a model of progressive MS ^48^. This model used non-obese diabetic mice, which have a genetic background of high innate immunity activation and aberrant activation of microglia that leads to an exacerbation of symptoms at the chronic phase ^49^. Indeed, microglia depletion in the chronic phase (not tested before) blocked EAE progression ^48^. Thus, the results of these studies are not comparable with ours because both the genetic background of the mice and the time window of the treatment are different.

More importantly, microglia depletion exacerbates demyelination and impairs remyelination in a neurotropic coronavirus infection model of MS ^50^. The authors showed a higher accumulation of damaged myelin deposits and debris in microglia depleted mice, that leads to impaired myelin repair and prolonged clinical disease. They propose that microglia functions could not be compensated by infiltrating macrophages. These results, although in a different MS model, are at odds with our data as we did not detect a deficit on myelin clearance, suggesting that in the EAE paradigm used in the current study macrophages compensate and efficiently phagocytose myelin. Accordingly, whereas microglia depletion increases monocyte infiltration in EAE lesions in the spinal cord, which could compensate for the microglia loss, it decreases the number of macrophages in the virus model of MS ^50^.

To sum up, we provide solid evidence showing that microglia and meningeal macrophages limit during EAE induction the infiltration and dispersion of peripheral macrophages throughout the CNS. Moreover, our results suggest that microglia are not essential for the development of chronic EAE nor for proper remyelination in this model. Moreover, we described a microglia-related mechanism promoting early antigen presentation in the CNS, by modulating the expression of B7 co-stimulatory molecules in other APCs, such as DCs or other myeloid populations. This lack of antigen presentation to infiltrating T cells delays the reactivation of these lymphoid cells and therefore provokes a slowdown of the EAE early events.

## ACKNOWLEDGMENTS

We kindly acknowledge the Microscopy Facilities of both the University of the Basque Country and Achucarro Basque Center for Neuroscience for the technical support. This work was supported by the Spanish Ministry of Economy and Competitiveness, (SAF2016-75292-R), Spanish Ministry of Science and Innovation Competitiveness (PID2019-109724RB-I00 and MCIN/AEI/10.13039/501100011033), Basque Government (IT1203/19, PIBA2016-1-0016 and PIBA-2020-1-0030), Centro de Investigación Biomédica en Red, Enfermedades Neurodegenerativas (CIBERNED) (grant no. CB06/05/0076) and ERDF “A way to make Europe” (RTI2018-099267-B-I00). AM, and AZ and NRI received predoctoral fellowships from the Spanish Ministry of Education and Science (AM) and the University of the Basque Country EHU/UPV (AZ, NRI), respectively.

## AUTHOR CONTRIBUTIONS

M.D. contributed to the conception and design of the study, data interpretation, and manuscript writing. A.M. contributed to the data acquisition and analysis, data interpretation, and manuscript writing. A.Z. contributed to data acquisition and analysis. M.E.L, B.R. and T.M. contributed to the analysis and interpretation of immune populations by FACS. N.R.L. contributed to the generation of the *Ccr2*^RFP/+^*fms*^EGFP/+^ mice colony. A.S. contributed to the generation of this colony, and to the revision of the manuscript. C.M. contributed to EAE experiments and to the revision of the manuscript. All authors have read and approved the submitted version, and agree to be personally accountable for their own contributions.

